# Tryptophan residues in TDP-43 and SOD1 mediate the cross-seeding and toxicity of SOD1

**DOI:** 10.1101/2020.07.27.224188

**Authors:** Edward Pokrishevsky, Michèle G. DuVal, Luke McAlary, Sarah Louadi, Silvia Pozzi, Andrei Roman, Steven S Plotkin, Anke Dijkstra, Jean-Pierre Julien, W. Ted Allison, Neil R. Cashman

**Affiliations:** Department of Medicine, Djavad Mowafaghian Centre for Brain Health, University of British Columbia, Vancouver, BC V6T 2B5, Canada; Department of Biological Sciences, Centre for Prions & Protein Folding Disease, University of Alberta, Edmonton, AB T6G 2E9, Canada; Department of Physics and Astronomy, University of British Columbia, Vancouver, BC V6T 1Z1, Canada; CERVO Brain Research Center, 2601 Chemin de la Canardière, Québec, QC G1J 2G3, Canada; Department of Pathology, Amsterdam Neuroscience, Amsterdam University Medical Centre, location VUmc, Amsterdam, The Netherlands

## Abstract

Amyotrophic lateral sclerosis (ALS) is a fatal neurodegenerative disease of motor neurons. Neuronal superoxide dismutase-1 (SOD1) inclusion bodies are characteristic of familial ALS with SOD1 mutations, while a hallmark of sporadic ALS is inclusions containing aggregated wild-type TAR DNA-binding protein 43 (TDP-43). Co-expression of mutant or wild-type TDP-43 with SOD1 leads to misfolding of endogenous SOD1 and aggregation of SOD1 reporter protein G85R-GFP in HEK293FT cells, and promotes synergistic axonopathy in zebrafish. This pathological interaction is dependent upon natively solvent-exposed tryptophans in SOD1 (tryptophan-32) and TDP-43 RRM1 (tryptophan-172), in concert with natively sequestered TDP-43 N-terminal domain tryptophan-68. TDP-43 RRM1 intrabodies reduce wild-type SOD1 misfolding in HEK293FT cells, via blocking tryptophan-172. Tryptophan-68 becomes antibody-accessible in aggregated TDP-43 in sporadic ALS motor neurons and cell culture. 5-fluorouridine inhibits TDP-43-induced G85R-GFP SOD1 aggregation in HEK293FT cells, and ameliorates axonopathy in zebrafish, via its interaction with SOD1 tryptophan-32. Collectively, our results establish a novel and potentially druggable tryptophan-mediated mechanism whereby two principal ALS disease effector proteins might directly interact in disease.

## Introduction

Amyotrophic lateral sclerosis (ALS) is characterized by progressive paralysis of the muscles of the limbs, speech, swallowing and respiration, usually leading to death within 2-5 years. Although the familial forms of the disease can be caused by mutation of ~20 genes, including Cu/Zn superoxide dismutase (SOD1) (1) and trans-activation response DNA-binding protein (*TARDBP*, encoding the protein TDP-43) (2), pathologically aggregated wild-type TDP-43 inclusions are a hallmark of all non-SOD1/fused in sarcoma (FUS)-associated ALS (3, 4). While mutations in SOD1 lead to robust intracellular aggregates of the protein in patient neurons, misfolded wild-type SOD1 can be detected by various SOD1 misfolding-specific antibodies in some sporadic ALS cases, in a granular and homogeneously distributed pattern (5–9). We have previously shown that transfection-mediated overexpression of mutant and wild-type TDP-43 induce the misfolding of endogenous human wild-type SOD1 in cultured cell lines, as well as in primary spinal cord cells derived from human wild-type SOD1 transgenic mice (5). The mechanisms whereby TDP-43 can induce misfolding of wild-type SOD1, and whether this process has significance for the pathobiology of human ALS, has been unknown.

We have found that the prion-like seeding and propagation of human wild-type SOD1 misfolding requires a tryptophan-tryptophan interaction in neighboring SOD1 molecules mediated by the single tryptophan at position 32 in SOD1 (Trp32) (10–12). Also, Trp32 mediates SOD1 aggregation in cultured cells, as well as zebrafish axonopathy *in vivo* (13, 14). Furthermore, drugs that interact with Trp32 (15, 16) can block propagated SOD1 aggregation in cell cultures and its toxicity *in vivo* (13, 14). Trp32 in SOD1 is estimated to be more solvent exposed than 90% of other tryptophans in the human structural proteome (10), an oddity consistent with pathological functionality, such as mediating intermolecular interactions between SOD1 and other molecules. Indeed, an aberrant interaction between, and co-aggregation of, mutant SOD1 with the stress-granule modulating protein G3BP1 depends on a three-way collaboration of Trp32 in SOD1 and a pair of aromatic residues (Phe380 and Phe382) in G3BP1 (17). Given the role of solvent exposed Trp in the misfolding and aggregation of SOD1, we speculated that cross-seeding of human SOD1 by TDP-43 might also be dependent on intermolecular interactions between solvent exposed Trp residues in both proteins. TDP-43 possesses six Trp residues, five of which are substantially solvent exposed in the native structure: two in the structured RNA recognition motif (RRM1) responsible for binding DNA and RNA (PDB: 4IUF (18), 4Y0F (19)), and three in the poorly structured low complexity C-terminal domain. The sixth tryptophan, Trp68, resides in the N-terminal domain (NTD), and is not solvent accessible in its natively folded structure. We now show that NTD Trp68 becomes antibody-accessible when aggregated in cultured cells and in sporadic ALS post-mortem spinal cord. We demonstrate that TDP-43-induced SOD1 misfolding, aggregation, and toxicity *in vivo* and *in vitro* are mediated by a three-way collaboration between two natively exposed tryptophans (SOD1 Trp32 and TDP-43 RRM1 Trp172), and the misfolding-exposed TDP-43 Trp68. These findings are further supported by blockade of wild-type SOD1 misfolding by novel intrabodies directed at TDP-43 RRM1, and by inhibition of cellular aggregation and zebrafish axonopathy by the Trp-interacting small molecule 5-fluorouridine. This pathological intermolecular interaction between tryptophan residues in TDP-43 and SOD1 may bridge a mechanistic gap in ALS pathogenesis, and may provide novel treatment targets for sporadic ALS.

## Results

### Interaction between tryptophan residues mediates the intermolecular cross-seeding of SOD1 by TDP-43 in cultured cells

We have previously reported that HEK293FT cells overexpressing either wild-type or mutant nuclear localization signal (ΔNLS) TDP-43 (TDP-43^ΔNLS^) can convert endogenous wild-type human SOD1 to a misfolded, propagating, and toxic form (5, 13). Since SOD1 seeding and misfolding propagation is dependent on its single tryptophan residue, Trp32 (10, 13, 14), we speculated that TDP-43 might induce SOD1 misfolding and aggregation via one or more of its six Trp residues (Fig 1A). Co-transfection of HEK293FT cells with SOD1^G85R^-GFP reporter (13, 20) and wild-type TDP-43 or nuclear localization signal mutant TDP-43^Δ NLS^ revealed that SOD1^G85R^-GFP aggregation was again strongly induced by these cytosolically mislocalizing and aggregating TDP-43 variants (Fig 1B), supported by live-cell kinetic analysis (Fig 1C). The induction of SOD1^G85R^-GFP aggregation by wild-type TDP-43 or TDP-43^ΔNLS^ was confirmed independently by flow cytometry analysis showing a significant 3-fold increase in the induction of SOD1^G85R^-GFP aggregation (Fig 1D and F). When all six of the Trp residues wild-type TDP-43 or TDP-43 ^ΔNLS^ were mutated to serine (tryptophan-less or Trpless), we found that co-expression of Trpless TDP-43^ΔNLS^ or Trpless wild-type TDP-43 with SOD1^G85R^-GFP resulted in a significant decrease of induced SOD1^G85R^-GFP aggregates forming across time as captured by time-lapse live-cell microscopy (Fig 1C and E). This decrease in the seeding of SOD1^G85R^-GFP was confirmed using flow cytometry (Fig 1D and F). Importantly, the Trpless TDP-43 variants were expressed at comparable abundance to TDP-43^Δ NLS^ (Fig 1B).

**Figure 1:**
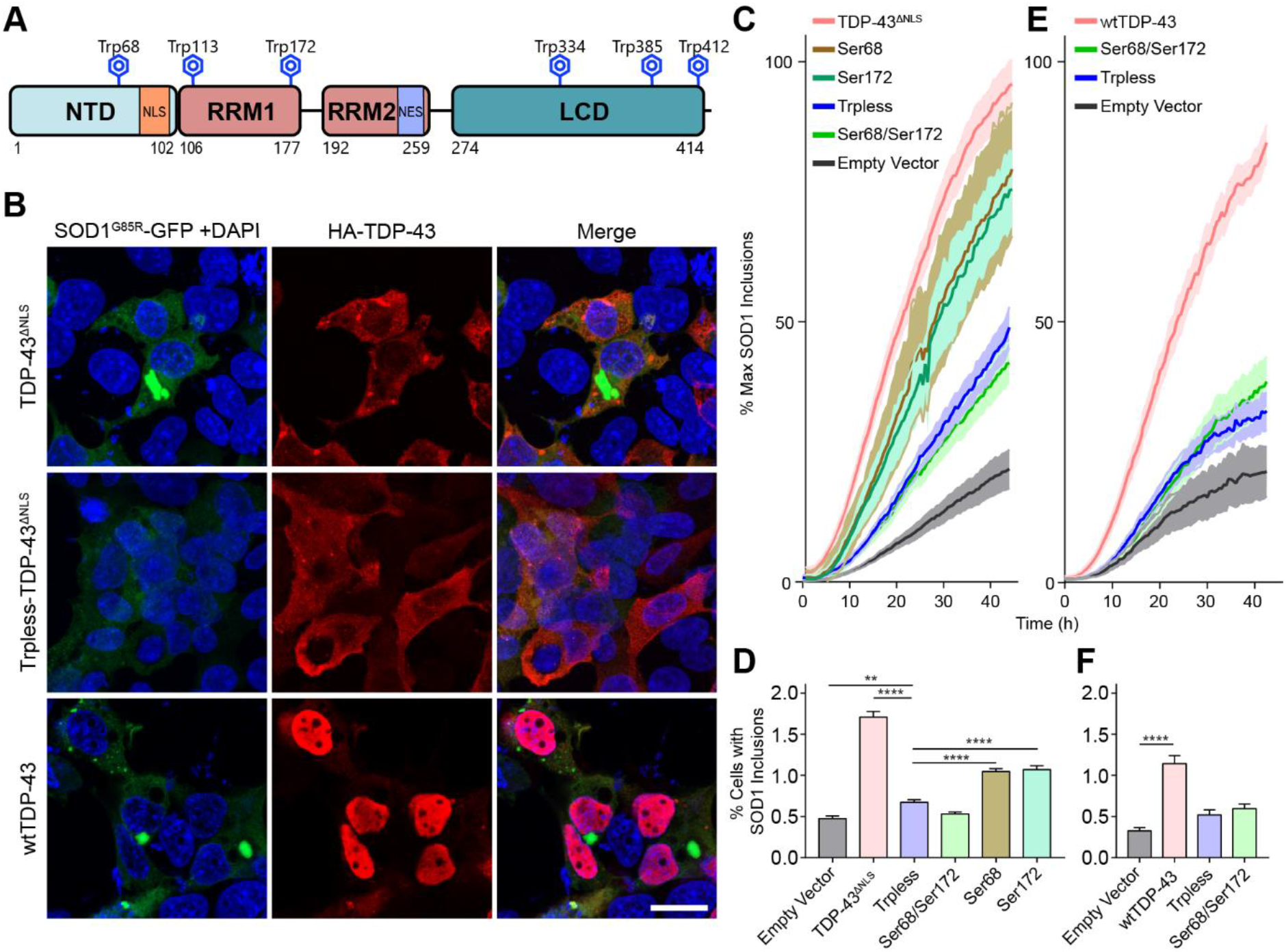
Tryptophan residues in wtTDP-43 and TDP-43^ΔNLS^ are required for the aggregation of mutant SOD1^G85R^-GFP in cultured cells. (**A**). Schematic representation of the TDP-43 protein highlighting its six Trp residues (yellow stars). NTD-N-terminal domain; NLS-nuclear localization signal RRM-RNA recognition motif; NES-nuclear export signal; LCD low-complexity domain. (**B**). TDP-43 variants (red) cross-seed aggregation of SOD^G85R^-GFP reporter protein (green), viewed 48 h after co-transfection into HEK293FT cells. TDP-43^ΔNLS^ with all Trp residues mutated to serine (Trpless-TDP-43^ΔNLS^) is abundantly expressed, but does not induce SOD1^G85R^-GFP aggregation. Scale bar: 20 μm. (**C**). Time-course analysis of induced SOD1^G85R^-GFP aggregate abundance in HEK293FT cells from 24 to 72 h post co-transfection with TDP-43^ΔNLS^, wherein combinations of Trp residues are mutated to serine (Ser). Trp68 and Trp172 in TDP-43 are required for induction of SOD1^G85R^-GFP aggregation. (**E)**. As per transfections of panel C except using wild-type TDP-43 plasmid variants. **D, F**. Percent of cells with detectable SOD1^G85R^-GFP aggregates quantified by flow cytometry shows more cross seeding of SOD1^G85R^-GFP by TDP-43^ΔNLS^ (**D**) or wild-type TDP-43 (**F**) constructs when compared to their variants bearing Trp to Ser mutation(s). Statistical significance was determined using one-way ANOVA and Dunnett’s test for multiple comparisons (**p < 0.01; ***p< 0.001; ****p< 0.0001). Error bars represent SEM; between 8 and 34 biological replicates were performed for each construct.

We next performed combinatorial step-wise Trp to Ser mutagenesis in order to identify which of the six Trp residues in TDP-43 are required for inducing SOD1^G85R^-GFP aggregation (Supp Fig 1). From this analysis, Trp68 and Trp172 were identified as crucial in SOD1^G85R^-GFP aggregation, with their combined mutation to Ser leading to a significant 2-fold decrease in the SOD1^G85R^-GFP aggregation rate (Fig 1C, D, Supp Fig 2A), and a 3-fold reduction in SOD1^G85R^-GFP aggregation measured via microscopy and flow cytometry (Fig 1E and F). Indeed, mutagenesis of both Trp68 and Trp172 reduced SOD1^G85R^-GFP aggregation to levels comparable to Trpless TDP-43^ΔNLS^, or to the vector-only negative control (reporter only; no TDP-43^ΔNLS^ in the co-transfection) (Fig 1C and D). Further Trp substitutions other than Trp68 and Trp172 did not significantly inhibit SOD1^G85R^-GFP aggregation, including mutation of Trp113 to produce a triple substitution of the Trp68 and the two RRM1 Trps. Trp to Ser mutations of the three C-terminal Trps in the TDP-43 low-complexity domain, which have been previously shown to participate in TDP-43 self-aggregation (21) did not affect SOD1^G85R^-GFP aggregation in co-transfected HEK293FT cells (Supp Fig 1). In sum, TDP-43, as wild-type and TDP-43^ΔNLS^ variants, are able to induce SOD1^G85R^-GFP aggregation in HEK293FT cells, by a mechanism that requires two collaborating N-terminal Trp residues Trp68 and Trp172.

### SOD1 and TDP-43 synergize to cause zebrafish motor axonopathy via tryptophan residues

Despite being an acute model, zebrafish axonopathy has proven to be an informative and tractable proxy for *in vivo* events associated with ALS etiology (22). Only wild-type TDP-43 and SOD1 could participate in sporadic ALS, so we assessed zebrafish primary motor axons for abnormal morphology in larvae injected with mRNA encoding human wild-type SOD1 Trp variants and/or human wild-type TDP-43 Trp variants. Overexpression of wild-type human SOD1 or TDP-43 in zebrafish has been previously found to engender abnormal axon branching (14, 22–26). Assessing axonopathy of primary motor neurons in the zebrafish ALS model allowed us to efficiently compare several Trp to Ser variants of SOD1 and TDP-43 *in vivo*, powered by robust sample sizes. Healthy primary motor axons extend from the spinal cord, past the notochord to innervate the trunk muscles (Fig 2A); these axons rarely exhibit branching above the ventral notochord boundary at 34-36 hpf, and abnormal axon morphology includes branching above this boundary (Fig 2A). Abnormal branching may be associated with muscle fiber patterning as well (Fig 2A). Overexpression of human wild-type SOD1 significantly elevated axonopathy by 1.3 fold over controls, similarly to previous reports (14, 25, 26), and expression of human wild-type TDP-43 in isolation increased axonopathy by 15% (Fig 2B). Compared to mRNA controls, the co-expression of SOD1 and TDP-43 significantly increased axonopathy by approximately 1.8 fold, the synergy of which is consistent with our hypothesis that TDP-43 induces SOD1 misfolding and toxicity (Fig 2B).

**Figure 2:**
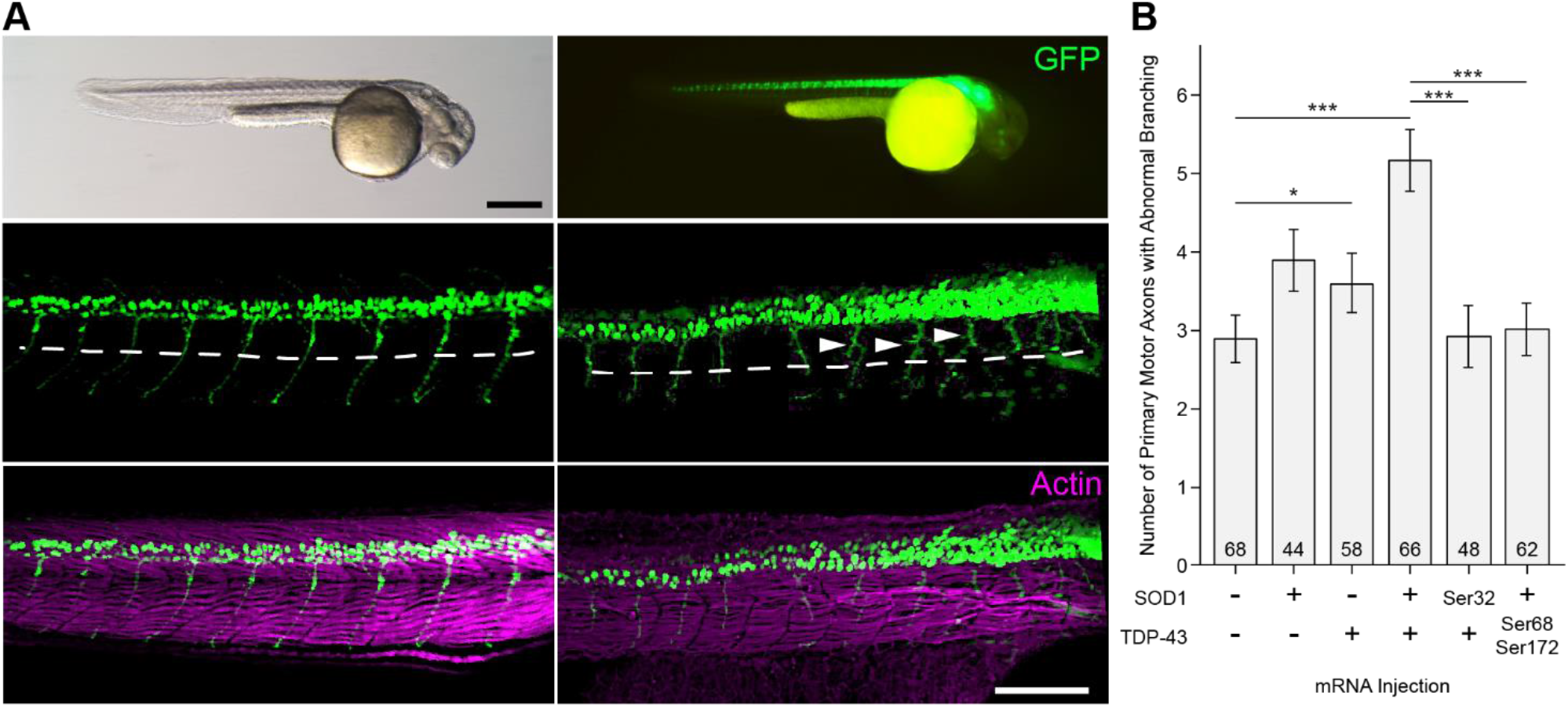
Tryptophan residues in human SOD1 and TDP-43 proteins mediate their synergistic impact on motor neuron pathology *in vivo*. (**A**). *mnx1:GFP* zebrafish (36 hpf, hours post-fertilization) expressing GFP in the motor neurons. Abnormal primary axons (arrowheads) with branches above the ventral notochord (dashed line). Muscle actin counterstained in magenta. Scale bars 0.5 mm (top left and top right); 200 μm (bottom 4 panels). (**B**). Human wild-type SOD1 + wild-type TDP-43 synergistically induce primary motor neuron axonopathy. Ser68 and Ser172 TDP-43 mutations reduced SOD1-TDP-43-induced axonopathy (p = 3.57 × 10^-5^). The SOD1 Trp32Ser mutation also abrogated axonopathy (p = 4.54 × 10^-5^), and in combination with Trpless-TDP-43^ΔNLS^ (p = 8.94 × 10^-5^). Error bars indicate SEM. * p < 0.05, ** p < 0.01, *** p < 0.001 (Kruskall-Wallis test with Mann-Whitney pairwise comparisons).

We next proceeded with substituting Trp residues in wild-type SOD1 and/or wild-type TDP-43 in combination to determine the impact of the Trps identified in the HEK293FT cell system detailed above. Substitution of Trp32 in SOD1 (SOD1^Trp32Ser^) abolished SOD1-TDP-43 induced axonopathy (p = 8.94 × 10^-5^) (Fig 2B), consistent with cell-based findings demonstrating that Trp32 is necessary for SOD1 to be induced to cross-seed by TDP-43. Again, we performed step-wise Trp substitutions in TDP-43 and found these to lead to statistically significant cumulative reductions in axonopathy, back to control mRNA levels. Mutation of Trp68 and Trp172 showed complete abrogation of the SOD1-TDP-43 synergistic impact (p = 3.57 × 10^-5^) (Fig 2C). These *in vivo* axonopathy results are consistent with the effect of Trp substitutions on SOD1 misfolding/aggregation as assessed in HEK293FT cells co-transfected with SOD1^G85R^-GFP and TDP-43 variants (Fig 1 and Supp Fig 2).

### Tryptophan 68 in TDP-43 NTD is exposed during misfolding/aggregation

TDP-43 RRM1 Trp172 appears to be solvent accessible on visual inspection of protein database (PDB) RRM1 example structures 4IUF (18) and 4Y0F (19) and Supp Fig 3A), and we have calculated that Trp172 solvent accessible surface area (SASA) is at the 91.8^th^ percentile compared to all Trps within a non-redundant PDB database of 27,015 proteins (27). However, TDP-43 Trp68 does not seem to be available for interaction at the TDP-43 molecular surface, and appears buried in the native structural core the NTD (Supp Fig 3B), at the 26.6^th^ percentile for SASA (27). We hypothesized that Trp68 acquires solvent exposure in misfolded/aggregated pathological TDP-43 in order to participate in the cross-seeding of wild-type SOD1. To test this hypothesis, we developed an affinity-purified rabbit polyclonal IgG generated against Trp68 in the context of its local amino acid sequence 65DAGWGNL71. Recombinantly expressed TDP-43 NTD demonstrated immunoreactivity with this antibody (named anti-Trp68) in a denaturing immunoblot system, but not in native gel immunoblots (Supp Fig 3C and D), demonstrating that loss of structure of NTD is required for antibody accessibility. Anti-Trp68 antibody also displayed reactivity to mislocalized cytoplasmic TDP-43 in transfected HEK293FT cells (TDP-43^ΔNLS^ and wild-type TDP-43; Fig 3A and Supp Fig 5). No immunoreactivity was detected in mock transfected and non-transfected HEK293FT cells (Supp Fig 4), showing the specificity of the anti-Trp68 antibody to misfolded/aggregated TDP-43 molecules. Interestingly, the anti-Trp68 antibody did not recognize TDP-43^ΔNLS^ cytoplasmic aggregates of a mutant Ser68 construct (Fig 3A), suggesting that immunoreactivity to this antibody is a good proxy of Trp-68 solvent exposure accompanying misfolding/aggregation. Anti-Trp68 displayed thread-like immunoreactivity against motor neurons in cervical spinal cord sections from two patients who died with sporadic ALS, in a similar pattern as a commercial anti-phosphorylated TDP-43 (Fig 3B), demonstrating that misfolding of the wild-type TDP-43 N-terminal domain occurs in ALS beyond the setting of artificial overexpression in cultured cells. Nuclei in unaffected neuronal and glial cells in these micrographs are not immunoreactive, consistent with the lack of Trp68 solvent exposure in normally folded healthy TDP-43. The above data support the notion that Trp68 is solvent exposed in pathological TDP-43 aggregates in cell culture *in vitro* and *in vivo* in human sporadic ALS, and is available in these conditions to collaborate with natively solvent-exposed Trp172 in RRM1 in the cross-seeding of SOD1 misfolding.

**Figure 3:**
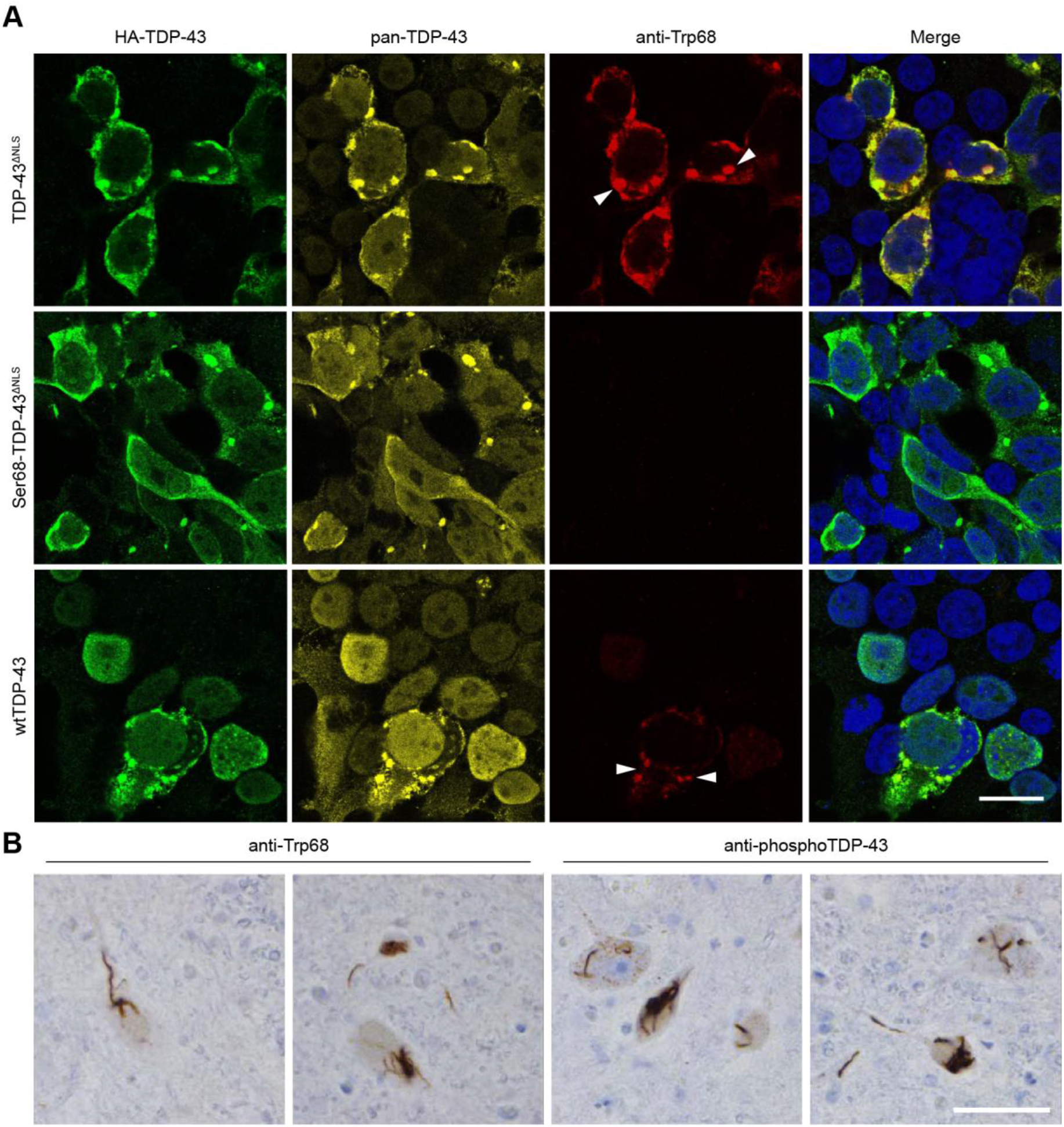
Trp68 is exposed in cytoplasmic TDP-43 aggregates in cultured cells and in sporadic ALS motor neurons. **(A).** The affinity-purified rabbit polyclonal anti-Trp68 antibody (red) was tested for reactivity and specificity in cells transfected with different TDP-43 constructs then fixed and stained 48 h later. A mouse pan-TDP-43 antibody against the C-terminal domain and a chicken anti-HA-tag antibody were used to test co-localization with TDP-43 (yellow) and HA-TDP-43 (green) respectively. The anti-Trp68 antibody specifically recognizes mislocalized cytoplasmic TDP-43 aggregates in TDP-43^ΔNLS^-transfected cells but not those lacking Trp68 (Ser68 TDP-43^ΔNLS^), nor does it recognize nuclear TDP-43. The composite images are a merge between nuclear, anti-Trp68, and anti-HA-tag staining. Scale bar: 20 μm. (**B**). Representative images of TDP-43 pathology in ALS cervical spinal cord sections from two subjects immunostained with anti-Trp68 in paraffin-embedded tissue. The left panel shows the anti-Trp68 antibody, and right the commercial pTDP-43 antibody. Motor neurons show thread-like inclusions using the anti-Trp68 antibody and the commercial pTDP-43 antibody, but no immunoreactivity is observed of natively folded TDP-43 in neuronal or glial nuclei Scale bar: 50 μm.

### The tryptophan-mediated cross-seeding activity of TDP-43 on SOD1 may constitute a target for ALS therapeutics

The apparent ability of aggregated TDP-43 to kindle misfolding of human wild-type SOD1, mediated via bi-molecular Trp residues as identified above, suggests that pathological TDP-43 possesses the potential to seed the propagated misfolding of human wild-type SOD1 (28). We have found that propagated misfolding of human wild-type SOD1 displays cytotoxicity *in vitro* (13), despite the fact that it does not form intracellular aggregates characteristic of mutant SOD1 familial ALS. If seeding of wild-type SOD1 by TDP-43 occurs in sporadic ALS, manifesting neurotoxicity in addition to TDP-43 toxicity mediated by nuclear depletion and cytoplasmic aggregation, then inhibiting the intermolecular interactions between TDP-43 and SOD1 may represent a target for interventions to slow disease progression. To test this idea, we employed two methods to interfere with the TDP-43-SOD1 cross-seeding in HEK293FT cells and in zebrafish. These methods target the interaction between TDP-43 and SOD1 via different tryptophans: 1) single-chain intrabodies directed against the RRM1 domain of TDP-43 (29), and 2) 5-fluorouridine, a small molecule that interacts with SOD1 at Trp32 (13–16, 20).

One of the key Trp residues in TDP-43 required for SOD1 aggregation, Trp172, is located in the RRM1 domain. We hypothesized that blocking this domain would attenuate the seeding of propagated misfolding of SOD1. We applied single chain antibodies (scFv) targeting the RRM1 domain of human TDP-43 (29). These antibodies were previously demonstrated to interact with cytoplasmic TDP-43, reducing its accumulation, and inhibiting NF-kB inflammatory signal generation (29). Here, we co-transfected HEK293FT cells with wild-type TDP-43 and intrabodies against the RRM1 (VH1Vk9 and VH7Vk9), and quantified SOD1 misfolding using 3H1 immunocytochemistry (5, 10). Consistent with our previous findings (5, 30), overexpression of wild-type TDP-43 triggered wild-type endogenous SOD1 misfolding (Fig 4A). SOD1 misfolding was significantly reduced in the presence of anti-RRM1 scFv ~2-fold compared to co-transfection of wild-type TDP-43 with empty-vector or control scFv (Fig 4B). Thus, RRM1 (including natively solvent-exposed Trp172) is likely key in TDP-43-induced kindling of SOD1 misfolding and propagation, and may constitute a target for therapeutic blocking of this pathological interaction.

**Figure 4:**
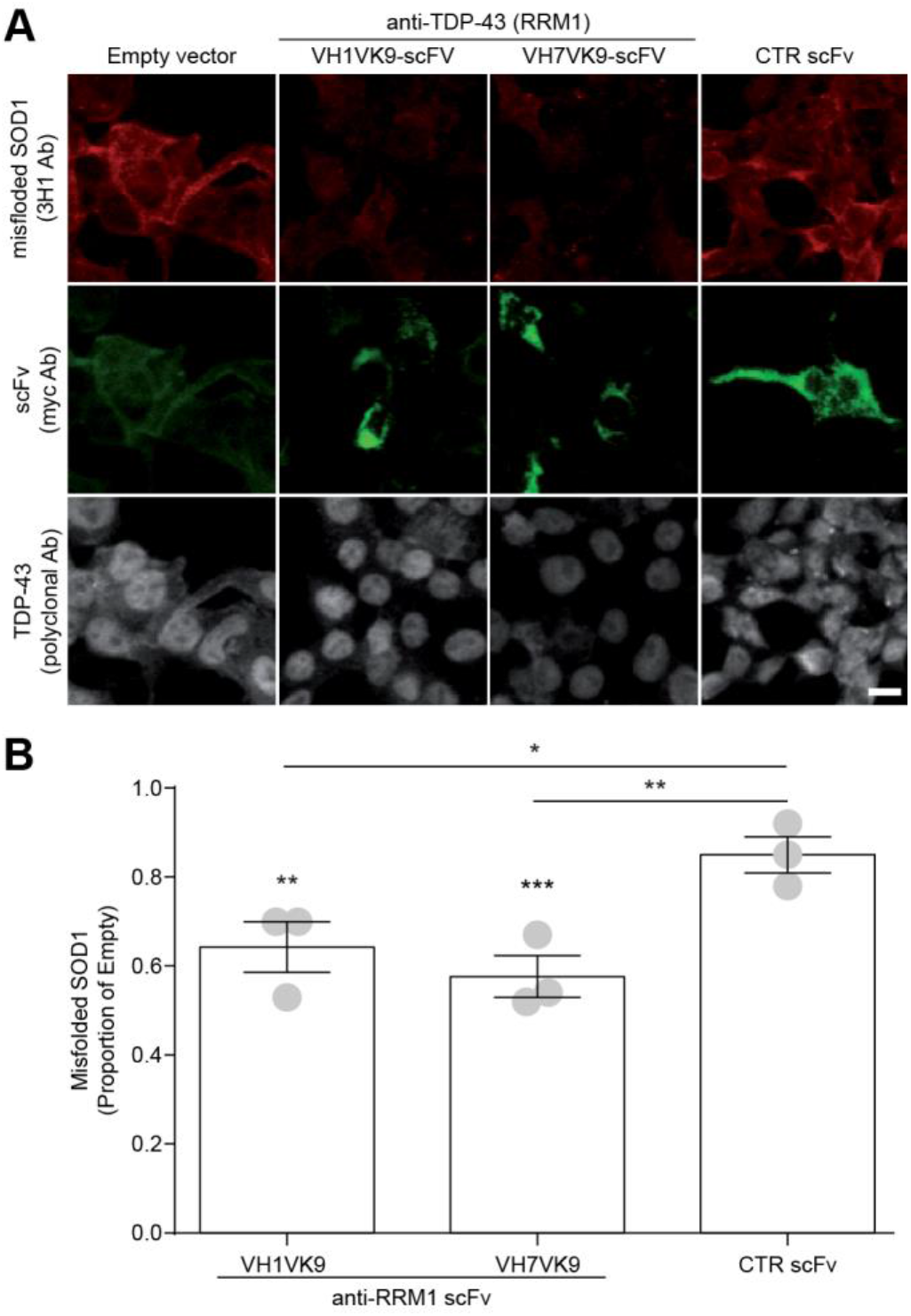
SOD1 misfolding in presence of overexpressed wild-type TDP-43 and anti-RRM1 single chain antibodies. (A) Representative picture of endogenous SOD1 misfolding (3H1 antibody, red) in HEK293 cells co-transfected with wild-type TDP-43 (gray) and anti-RRM1 scFvs (green) (VH1Vk9 and VH7Vk9), control scFv (D1.3, anti-chicken lysozyme) or empty plasmid for scFv. Scale bar = 10μm. (B) Quantification of misfolded SOD1. Statistical significance was established using one-way ANOVA followed by Tukey’s multiple comparison test, * = p < 0.05, **=p < 0.01 and *** = p < 0.001 versus Empty or versus control scFv antibody when shown by comparisons bars. Data are expressed as mean ± SEM of 3H1 integrated density normalized on number of cells from n = 3 different experiments (dots in the bar graph).

The human SOD1 Trp32 is required for the seeding and propagated misfolding of wild-type and mutant SOD1 (13, 14), and may thus represent another ALS therapeutic target. Because 5-fluorouridine (5-FUrd) can prevent induction of SOD1 aggregation by mutant SOD1 via blocking Trp32 (13–15), we speculated that this drug might also block TDP-43-induced propagated seeding of SOD1 reporter protein in cells, and alleviate axonopathy in zebrafish. 5-FUrd (1-5 μM) significantly inhibited the TDP-43 ^ΔNLS^-induced aggregation of SOD1^G85R^-GFP in HEK293FT cells (Fig 5A). 5-FUrd did not affect the aggregation status of TDP-43^ΔNLS^, suggesting that 5-FUrd inhibition is specific for SOD1 Trp32 sequence/conformation local site, which is not recapitulated in the local structure of TDP-43 Trps. Time-lapse live-cell microscopy revealed that the TDP-43-induced aggregation of SOD1^G85R^-GFP reporter protein is reduced proportionately with increasing concentrations of 5-FUrd (1 and 5 μM, Fig. 5B). The rate of SOD1^G85R^-GFP aggregation during the linear aggregate growth-phase (approximately 8-24 h after data acquisition or 24-40 h post co-transfection) apparently decreases with increasing concentrations of 5-FUrd; however, the correlation was not statistically significant (Supp Fig 5), possibly due to similar outcomes for both drug doses. The efficacy of 5-FUrd in reducing TDP-43-induced SOD1 aggregation was validated independently by quantifying cells with flow cytometry analysis (Fig 5C). A treatment of 5 μM 5-FUrd reduced SOD1^G85R^-GFP aggregation to levels comparable to the control, in which empty vector was used in place of TDP-43^ΔNLS^. These results support the notion that drugs targeting Trp32 may ameliorate SOD1-mediated neurotoxicity in ALS. Treatment of cells with the non-fluorinated pyrimidine base uridine did not inhibit aggregate formation, even at a concentration of 50 μM (Fig 5C-D). Finally, towards *in vivo* drug application, we considered that the toxicity of 5-FUrd could be mitigated during chemotherapy regimens by co-administering uridine (31). We confirmed that supplementing with uridine does not decrease the ability of 5-FUrd to reduce TDP-43-induced aggregation of SOD1 *in vitro* (Supp Fig 6).

**Figure 5:**
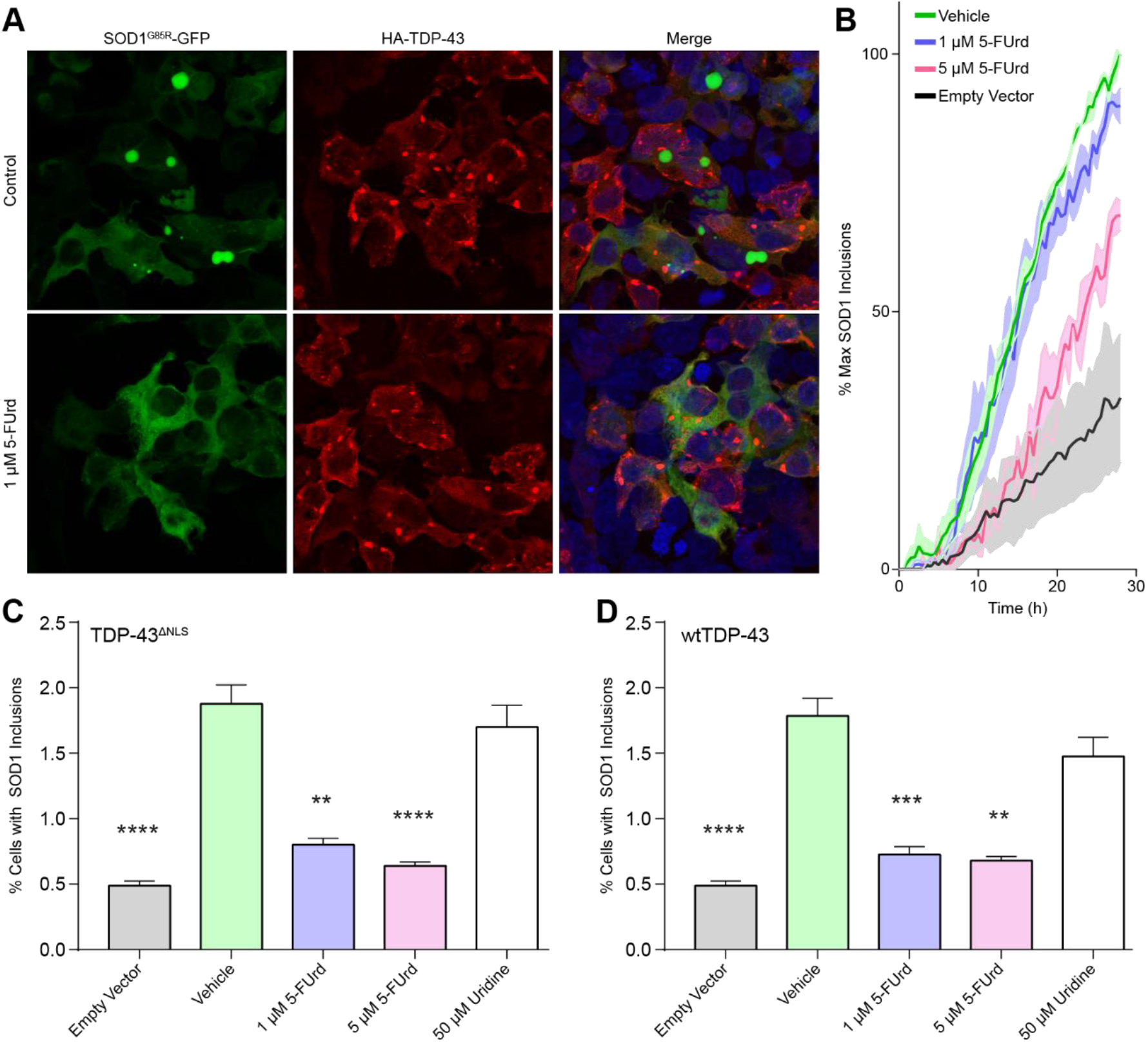
5-FUrd effectively blocks the cross-seeding of SOD1 by TDP-43. **A.** Immunocytochemistry of HEK293FT cells 48 h post co-transfection with TDP-43^ΔNLS^ (red) and SOD^G85R^-GFP reporter protein (green) in the presence of vehicle (top) or 1 μM 5-FUrd (bottom) demonstrating a reduction in SOD1 aggregates with 5-FUrd. Scale bar: 50 μm. (**B**). Time-lapse live-cell microscopy of SOD^G85R^-GFP cross-seeding by TDP-43^ΔNLS^ in the presence of 1 or 5 μM 5-FUrd shows a reduction in aggregate accumulation proportionate to drug concentration. Images were acquired every 30 minutes for 30 h. **C-D**. Flow cytometry of saponin-treated cells co-transfected with the reporter protein with either TDP-43^ΔNLS^ (**C**) or wild-type TDP-43 (**D**) showing a reduced number of SOD1^G85R^-GFP aggregate containing cells when treated with 5-FUrd. For each experiment, error bars represent SEM of 8-13 biological replicates. Statistical significance was determined using one-way ANOVA and Dunnett’s test for multiple comparisons (**p < 0.01; ****p < 0.0001).

Given our findings in HEK293FT cells, we next tested the capacity of 5-FUrd to rescue SOD1-TDP-43-induced axonopathy in zebrafish. A low dose of 1.5 μM 5-FUrd (+ 5 μM uridine) rescued the axonopathy phenotype significantly in SOD1 + wild-type TDP-43 injected embryos by 36%, and in SOD1 + TDP-43^ΔNLS^ embryos by 46% (Fig 6B and C).

**Figure 6:**
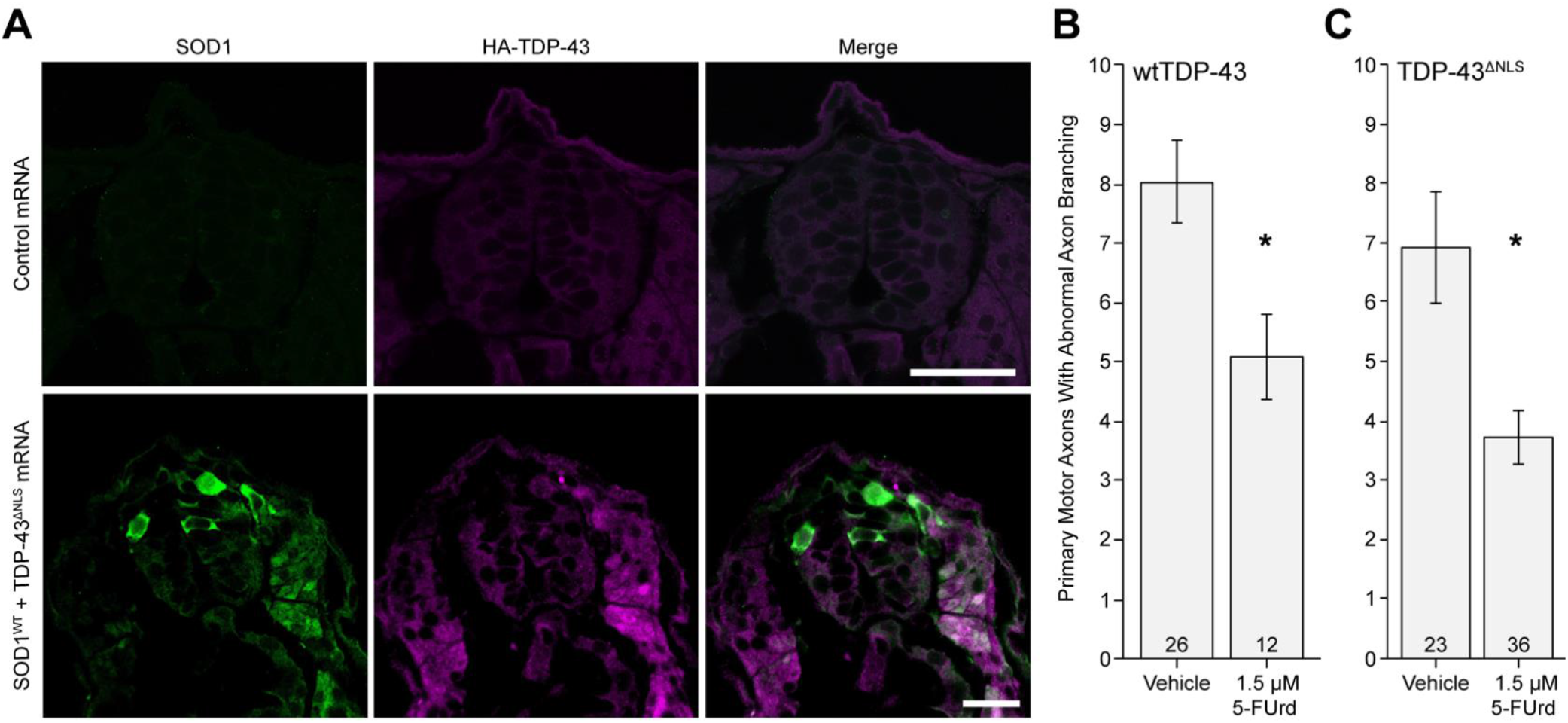
5-FUrd rescues axonopathy induced by co-expressing human SOD1 and wild-type TDP-43 in zebrafish. When human SOD1 and TDP-43 are co-expressed in zebrafish (**A**, exemplar cross-sections of spinal cord), the motor neurons exhibit axonopathy (**B,C**) and this axonopathy can be partially but significantly rescued with drugs that block human SOD1 Trp32. **(B)** 5-fluorouridine (5-FUrd, 1.5 μM) rescued axonopathy in embryos injected with SOD1 + wild-type TDP-43 (p = 0.011). (**C**) A significant reduction was also observed with 5-FUrd in SOD1 + TDP-43^ΔNLS^-injected embryos (p = 0.040). Both vehicle and 5-FUrd solutions contained 5 μM uridine and 0.2% DMSO. Error bars indicate SEM; * p < 0.05 (Mann-Whitney pairwise comparisons); sample sizes are noted at the base of each bar.

## Discussion

Whilst the formation of inclusions incorporating wild-type TDP-43 is a hallmark of sporadic ALS (4), several studies have identified the presence of misfolded, although typically unaggregated, wild-type SOD1 in TDP-43 sporadic ALS and non-SOD1 FALS using an array of SOD1 misfolding-specific antibodies (5–9, 32, 33). Misfolded, but unaggregated, endogenous wild-type SOD1 has also been detected in cell cultures overexpressing wild-type or cytosolically mislocalized mutant TDP-43 (5), which we have replicated in the current manuscript (Fig 4) in the course of testing TDP-43 RRM1 intrabodies. Once misfolded, this species of wild-type SOD1 (perhaps oligomeric; (34, 35)) can be transmitted to other cells, triggering additional rounds of induced misfolding of the endogenous wild-type SOD1 in recipient cells, and accompanied by cytotoxicity (30). Previous investigations have yielded little genetic interaction between SOD1 and TDP-43, including a failure of SOD1 mRNA to rescue TDP-43 knockdown- or mutant-induced axonopathy in zebrafish (36–41). Here, we tested a novel hypothesis in cell culture and zebrafish to demonstrate a pathologic synergy between SOD1 and TDP-43 *in vitro* and *in vivo*, which occurs at a post-translational level and is mediated by specific solvent-exposed Trp residues in both proteins. We find that overexpression of either mutant TDP-43^ΔNLS^ or wild-type TDP-43 can cross-seed SOD1^G85R^-GFP reporter protein in HEK293FT cell cultures, and that expression of either human wild-type TDP-43 or human wild-type SOD1 result in motor axonal toxicity in zebrafish, an effect amplified by co-expression of these two proteins.

As in human sporadic ALS, zebrafish TDP-43 toxicity is associated with cytosolic TDP-43, but misfolded human wild-type SOD1 immunoreactivity is not concentrated into discrete aggregates (Fig 6A). We also show that induced conversion of SOD1 species by TDP-43, in both HEK293FT cells and zebrafish, is specified by specific tryptophan residues in both proteins, militating for a direct proteinprotein interaction. For SOD1, the aggregation of SOD1^G85R^-GFP (in cultured cells) and cross-seeding of wild-type SOD1 (in zebrafish) is dependent on surface-exposed SOD1 Trp32, similar to earlier studies of SOD1-SOD1 propagated misfolding (10, 11, 13, 14, 20). Cross-seeding of wild-type SOD1 by TDP-43^ΔNLS^ and wild-type TDP-43, and SOD1^G85R^-GFP aggregation, are all dependent on the natively solvent-exposed RRM1 Trp172, and the misfolding/aggregation-induced exposure of Trp68 by loss-of-structure of the TDP-43 N-terminal domain. Interestingly, although we have found that misfolding/aggregation cytosolic exposure of TDP-43 Trp68 is a requirement for efficient cross-seeding of human SOD1, efficient conversion also requires the collaboration of one additional RRM1 Trp which is natively solvent-exposed. Moreover, the three solvent-exposed Trps in the C-terminal low complexity domain do not substitute for exposure of Trp68, indicating that the N-terminal structures NTD and RRM1 play a key role in this process which is driven by specific solvent-exposed Trps.

Pathological activities of mutant or wild-type TDP-43 are generally thought to include at least two distinct toxicities: loss-of-function of native TDP-43 that is accompanied by nuclear depletion, which includes defective RNA splicing, transport and stabilization, and DNA repair (42, 43), and gain-of-function from TDP-43 misfolding and cytosolic aggregation (44). These two toxicities may actually be synergistic, including acceleration of nuclear depletion from the recruitment of functional TDP-43 to the cytosolic aggregates by a prion-like mechanism (45). Although misfolded TDP-43 has well-delineated toxicity on its own, including the initiation of ER stress and mitochondrial dysfunction (46, 47), a recently recognized toxic activity of misfolded or aggregated TDP-43 is to trigger the misfolding and dysfunction of other proteins, including nuclear pore proteins and karyopherins (48), proteins involved in mRNA translation (RACK1, DISC1) (49, 50), and proteins implicated in other neurodegenerative diseases (tau, alpha synuclein) (51, 52). Thus, it is perhaps not surprising that SOD1 can be included in the “pathological interactome” of TDP-43.

Although the toxic prion-like propagation of mutant SOD1 is well recognized, human wild-type SOD1 may be an important substrate of TDP-43 seeding, because human wild-type SOD1 misfolding can propagate within cells and between cells *in vitro* (10, 53), potentially resulting in gain-of-function cytotoxicity (30) and/or loss-of-function from diminished dismutase activity (54). Recent data has implicated a role for toxic forms of wild-type SOD1 secreted from sporadic ALS astrocytes (55) and detected in sporadic ALS CSF (56). Interestingly, it has been reported that misfolded mutant SOD1 acquires its own pathological interactome, including G3BP1, α-synuclein, and amyloid-β (17, 57, 58). The concept that SOD1 misfolding and toxicity is downstream of TDP-43 is consistent with reports that the emergence of the sporadic ALS syndrome may be preceded by six mathematically defined preclinical “steps” (59), but that FALS mutations in TDP-43 are best captured by a model comprising four steps, and that SOD1 mutations apparently require only two steps (60), suggesting that wild-type SOD1 may not only express toxicity and/or loss-of-function in sporadic ALS, but that it may be downstream of other events in ALS pathogenesis, such as TDP-43 aggregation.

If misfolded wild-type SOD1 possesses toxic activities beyond those inherent in TDP-43 aggregates, the relevance of the identified set of Trp residues to the development of SOD1 and TDP-43 toxicity at the whole organism level is a compelling avenue for further exploration. We explored the therapeutic relevance of Trp32 in this TDP-43-SOD1 paradigm using 5-FUrd, a candidate drug predicted to block Trp32 at the molecular surface of human SOD1 protein (13–15, 61). Since 5-FUrd successfully reduces SOD1 aggregation in cell culture and reduces motor neuron toxicity in zebrafish, it may be an attractive candidate for further investigation, not only in mutant SOD1-familial ALS, but sporadic ALS as well. It is also possible that partial knockdown of human wild-type SOD1 through antisense oligonucleotides or RNA-based therapies might be tested in sporadic ALS, beyond the SOD1-mutant familial ALS for which these gene therapy modalities have recently demonstrated efficacy (62, 63).

## Contributions

**Conceptualization:** NRC, WTA, EP, MGD; **Methodology:** NRC, WTA, SSP, JPJ, EP, MGD, LM, SL, SP, AR, AD; **Validation:** NRC, WTA, EP, MGD.; **Formal analysis:** EP, MGD, LM, SL, SP, AR, SSP; **Investigation:** EP, MGD, SL, SP, AR, SSP; **Resources:** NRC, WTA, SSP, JPJ; **Data curation**: EP, MGD, LM, SL, SP, AR, SSP; **Writing - original draft:** NRC, WTA, EP, MGD; **Writing - review & editing**: NRC, WTA, SSP, JPJ, EP, MGD, LM, SL, SP; **Visualization:** EP, MGD, LM, SL, SP; **Supervision**: NRC, WTA, SSP, JPJ; **Project administration:** NRC, WTA.; **Funding acquisition:** NRC, WTA, SSP, JPJ.

## Acknowledgements

MGD was funded by CIHR and Alberta Innovates Health Solutions MD/PhD awards. SP received the 2019 Marlene Reimer Brainstar of the year award from CIHR-CAN (ICT-171454). SSP acknowledges support from CIHR Transitional Operating Grant 2682. JPJ acknowledges CIHR funding. Operating funds to WTA were from anonymous donors. NRC acknowledges grant support from ALS-Canada, Brain Canada, the R. Howard Webster Foundation, and the Canadian Consortium for Neurodegeneration in Aging. NRC also gratefully acknowledges generous donations from John Tognetti and William Lambert.

## Supplementary Information

**Supplementary Figure 1:**
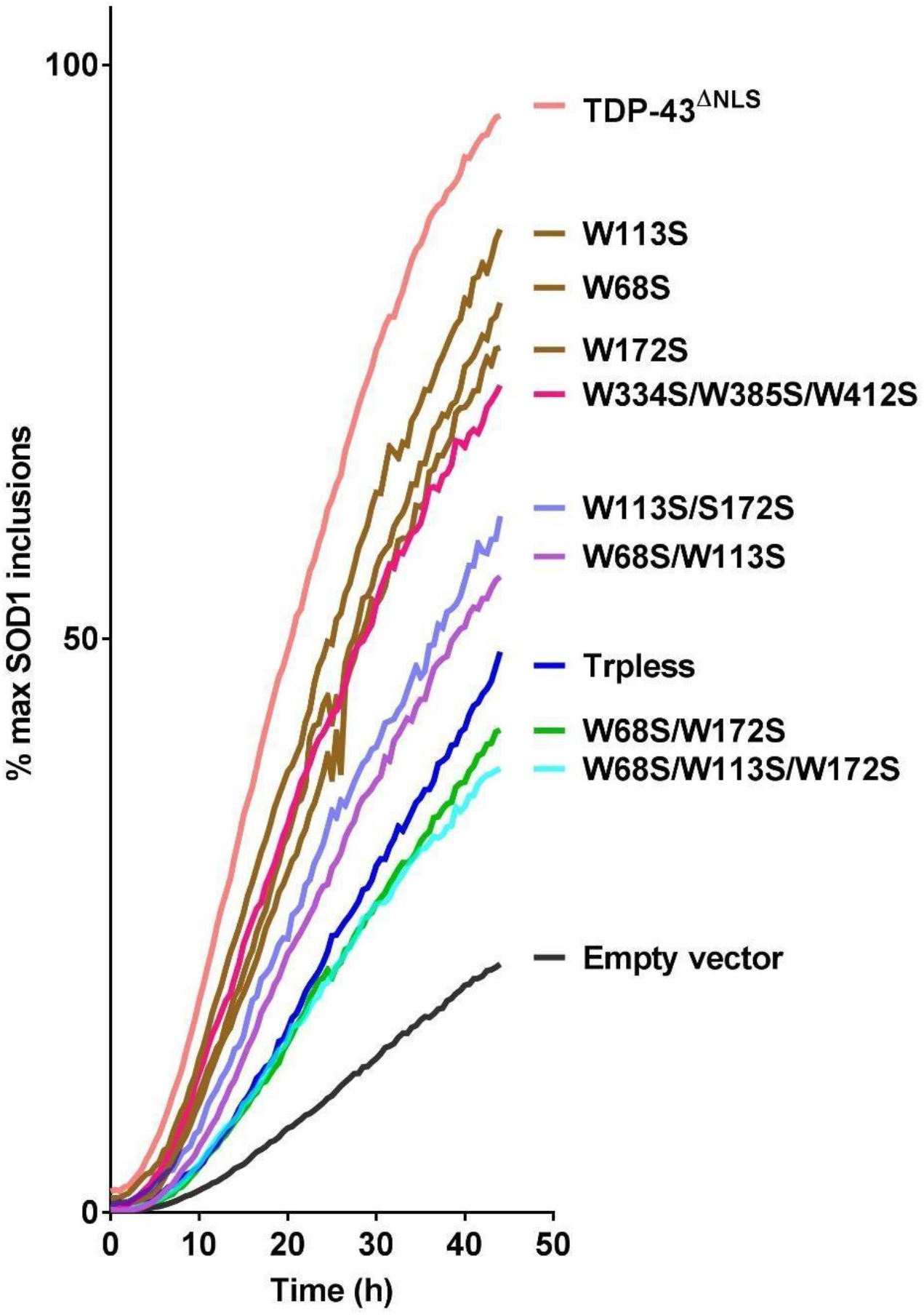
Tryptophan residues at position 68 and 172 in TDP-43^ΔNLS^ are most critical for aggregation of mutant SOD1-based reporter protein. Time-course algorithm count of induced SOD1 aggregates in HEK293FT cells from 24 to 72 h post co-transfection with the indicated TDP-43^ΔNLS^ and its variants. Images were acquired every 30 minutes. Time-point h = 0 corresponds to the beginning of imaging, which occurred approximately 16 h post-transfection. The number of inclusions at every time point is expressed as a percentage of the final inclusions of TDP-43^ΔNLS^ in the biological repeat. Error bars were removed for clarity reasons. Each curve represents 3-18 independent biological repeats.

**Supplementary Figure 2:**
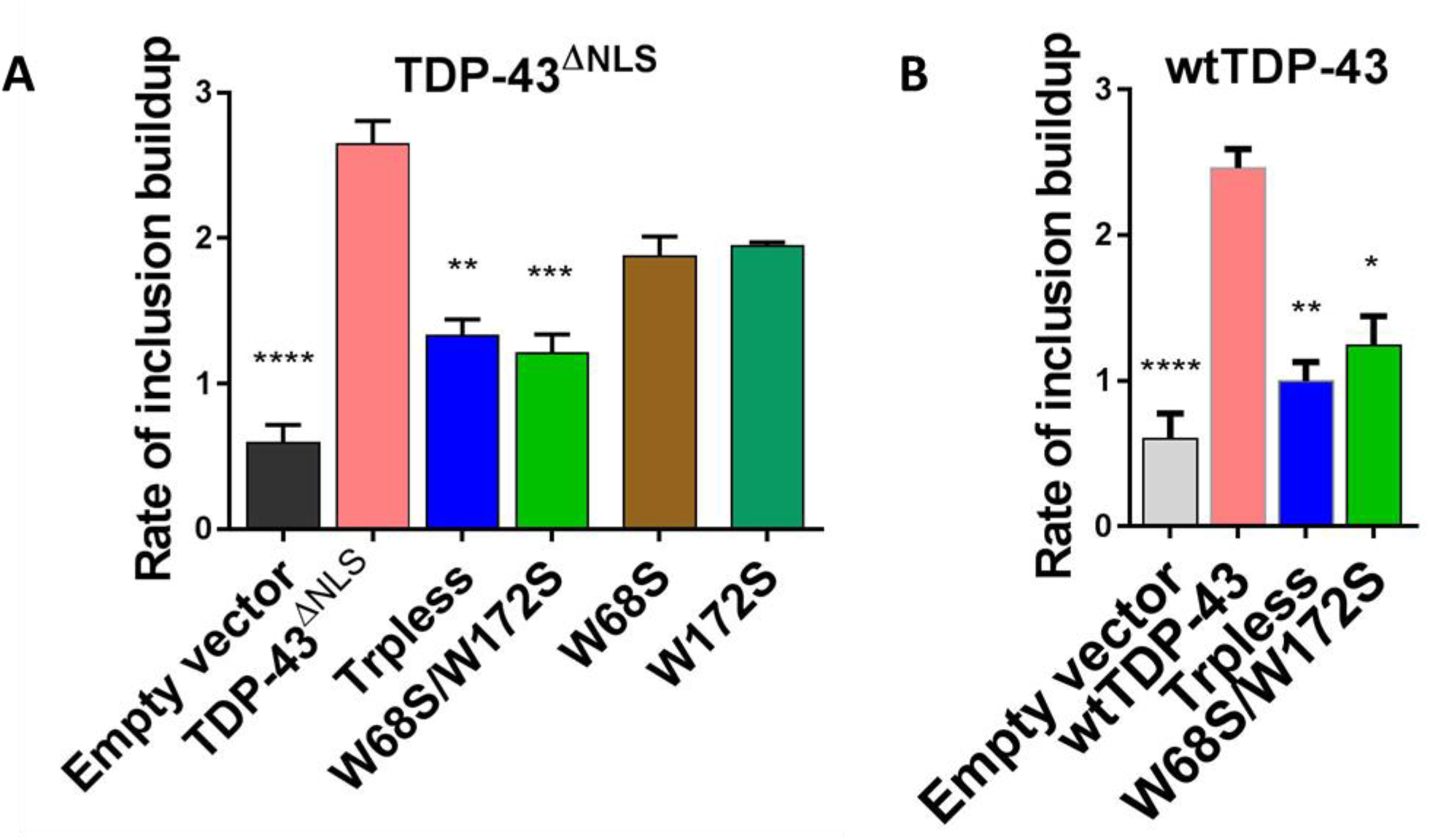
The rate of SOD1 inclusion formation is dependent on the presence of tryptophan residues in TDP-43. Cells co-transfected with SOD1 and TDP-43 show a rapid linear growth of SOD1 inclusions, whereas those with TDP-43 variants lacking all (Trpless) or two key tryptophans (Trp68Ser/Trp172Ser) exhibited a much slower linear growth. The rate of inclusion buildup per hour, plotted here, is based on the linear growth phase of the SOD1 inclusions in the presence of TDP-43^ΔNLS^ (**A**) or wild-type TDP-43 (**B**) based constructs and was quantified from 10-35 h after data acquisition or 26-51 h post co-transfection. Statistical significance was determined using one-way ANOVA and Dunnett’s test for multiple comparisons (* p < 0.05; ** p < 0.01; *** p < 0.001; **** p < 0.0001).

**Supplementary Figure 3:**
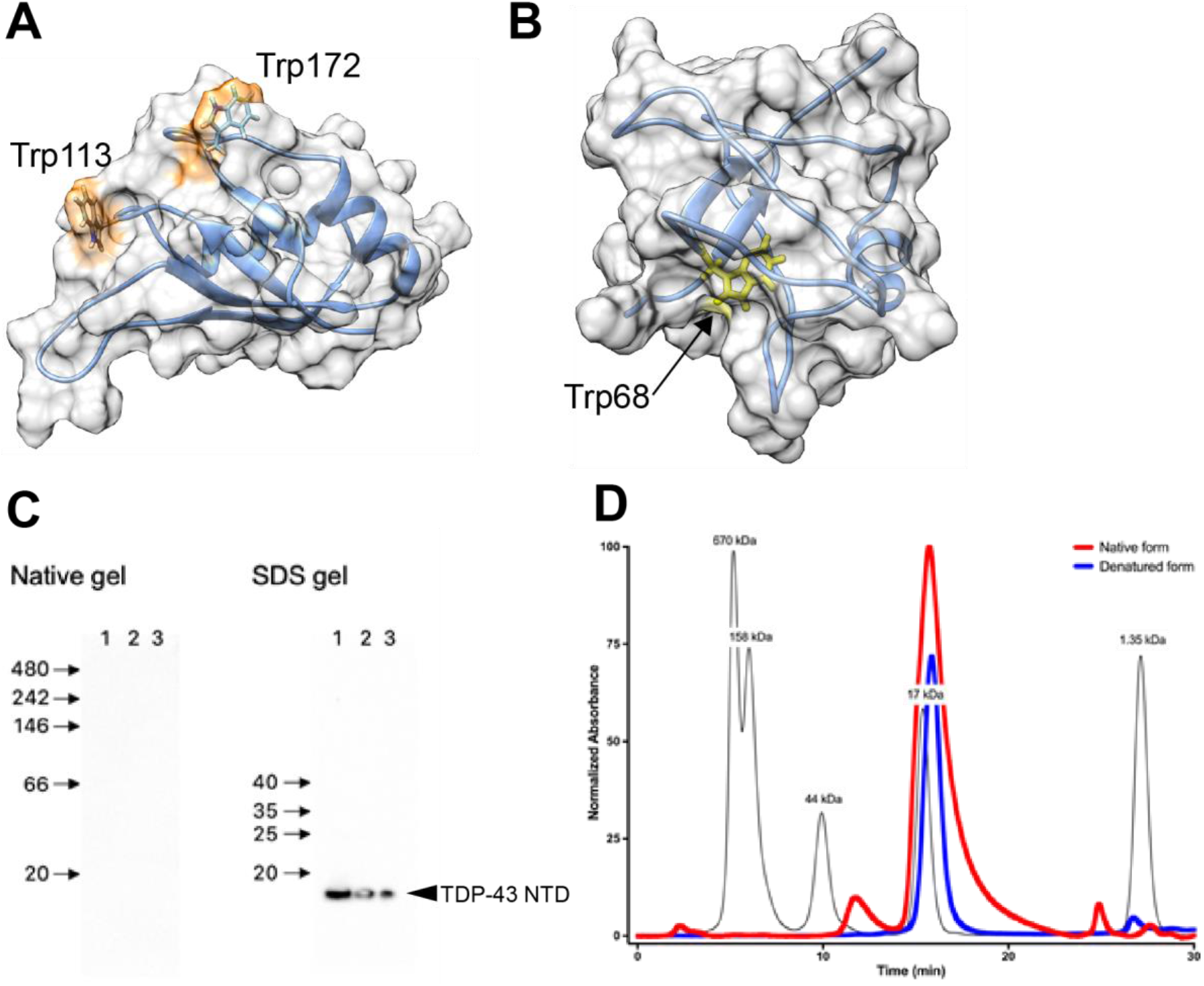
Analysis of the solvent accessible surface area (SASA) of tryptophan residues in the N-terminus of TDP-43. **(A)** Representative centroid structure of the RRM1 domain of TDP-43, obtained from equilibrium molecular dynamics simulation. Trps 113 and 172 are significantly exposed compared to most native structures in the protein databank (27). **(B)** Representative centroid structure of the N-terminal domain of TDP-43, obtained from equilibrium molecular dynamics simulation. Trp68 is nearly fully buried: only about 9% of its tripeptide Gly-Trp-Gly value of 247 Angstrom^2^ is exposed in the centroid structure. **(C)** The anti-Trp68 antibody used in immunoblot analysis of native- and SDS-PAGE gels shows reactivity only to denatured TDP-43 NTD. Lanes 1, 2, 3 contained 0.6, 0.3 and 0.15 μg protein per lane, respectively. **(D)** SEC fractionation chromatogram of TDP-43 NTD in native form (red line) and denatured in 6 M guanidinium chloride solution (blue line). The protein remains monomeric in both conditions. MW markers are superimposed for reference (black line).

**Supplementary Figure 4:**
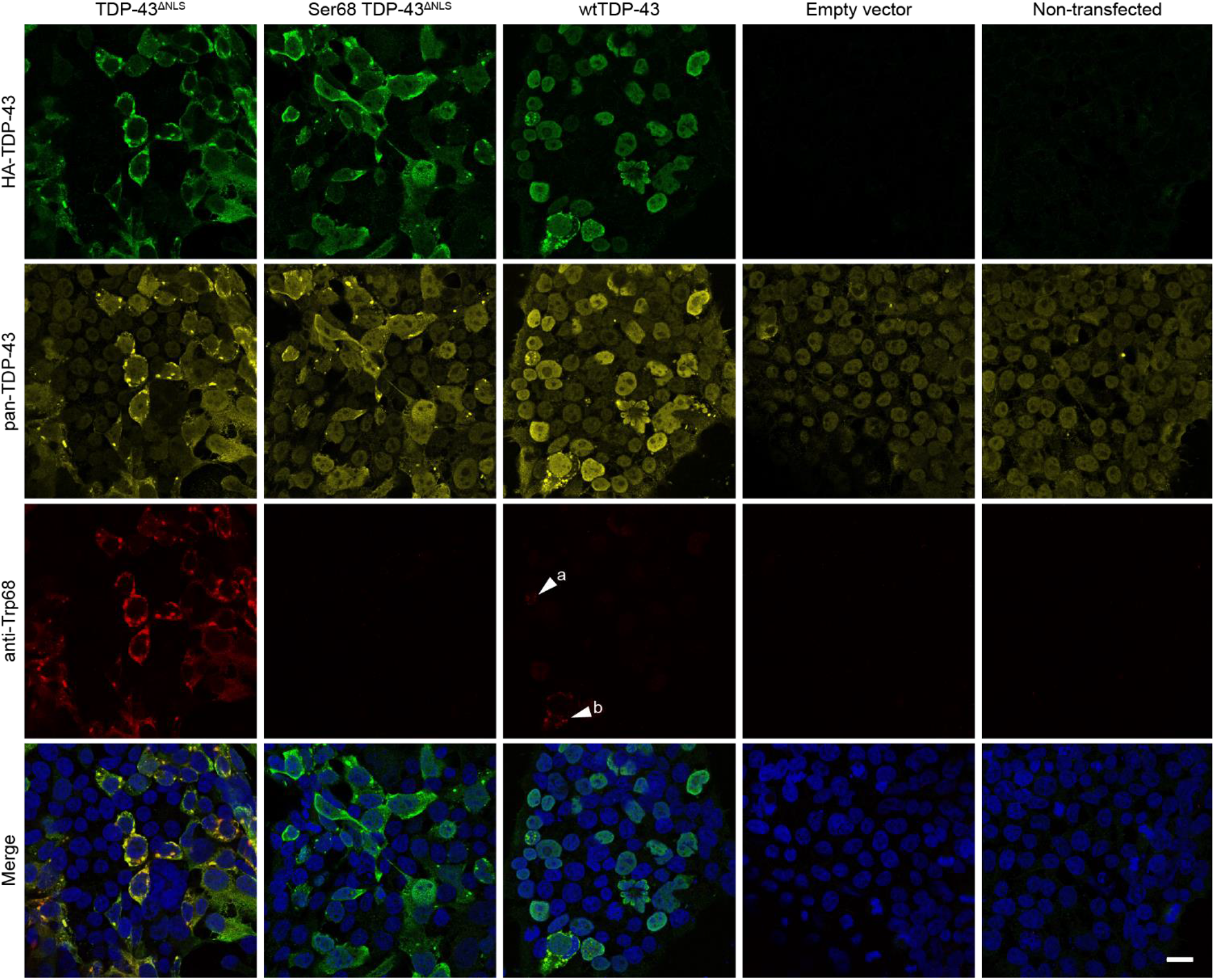
Trp68 is exposed in aberrant TDP-43-containing cytoplasmic and nuclear aggregates. The rabbit anti-Trp68 antibody (red) was tested for reactivity and specificity in cells transfected with different TDP-43 constructs. A mouse pan-TDP-43 antibody against the C-terminal domain (Proteintech, USA) and a chicken anti-HA-tag antibody (Abcam, USA) were used to test co-localization with TDP-43 (yellow) and over-expressed TDP-43 (green) respectively. The anti-Trp68 antibody specifically recognizes mislocalized cytoplasmic TDP-43 aggregates in TDP-43^ΔNLS^-transfected cells aberrant nuclear TDP-43 aggregates that form when TDP-43 is overexpressed. Trp68 is also exposed in cytoplasmic aggregates that form when TDP-43 is overexpressed in wild-type TDP-43 transfected cells, and these aggregates are recognized by the specific antibody. However, Trp68 does not stain non-aggregated nuclear TDP-43 nor does it recognize cytoplasmic TDP-43 aggregates lacking Trp68. No background staining was seen in mock transfected and non-transfected cells. Scale bar: 20μm.

**Supplementary Figure 5:**
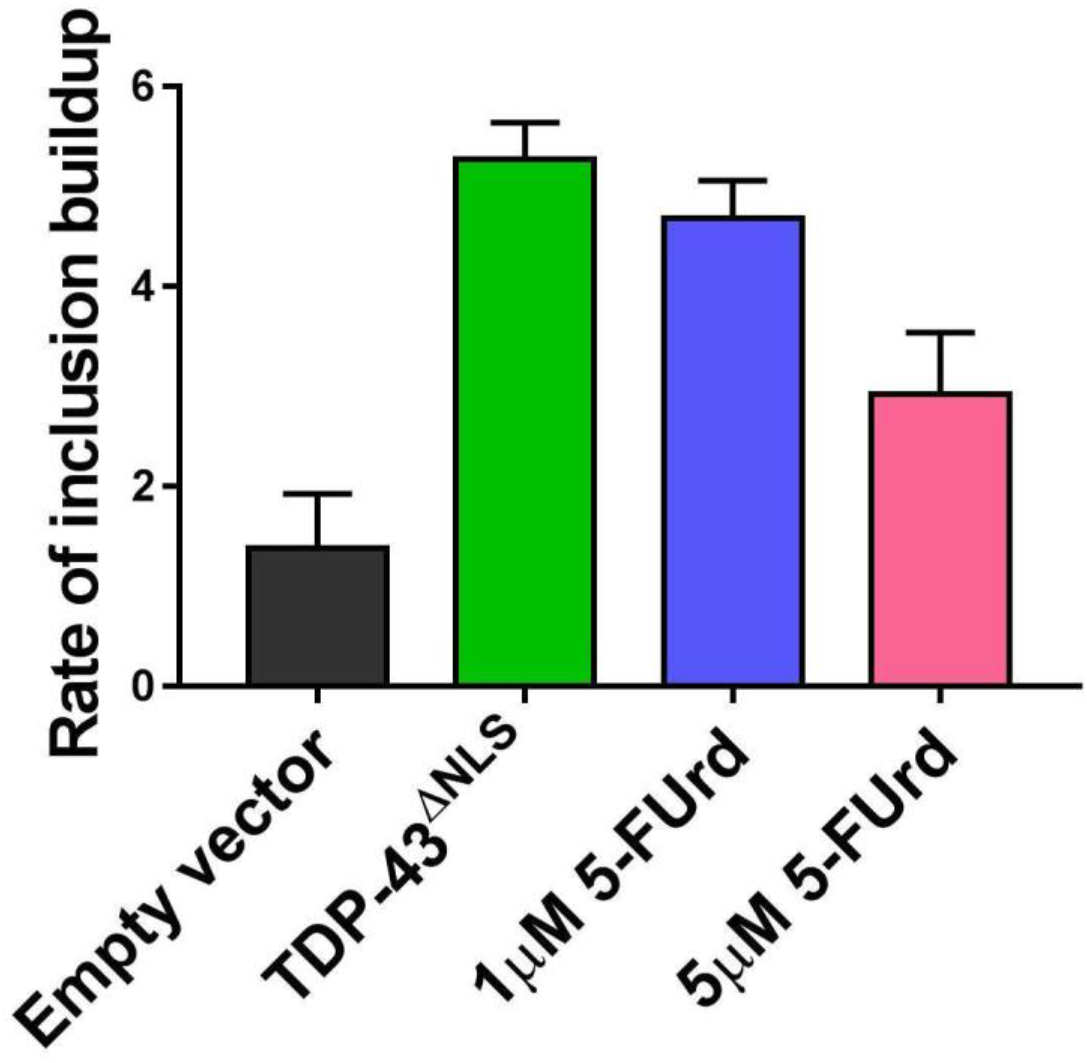
The rate of TDP-43-induced SOD1 inclusion formation is decreased in the presence of 5-FUrd. The rate of inclusion build-up per hour is based on the linear growth phase (approximately 8-24 h after data acquisition or 24-40 h post co-transfection) of TDP-43-induced SOD1 aggregation in the presence of 1 or 5 μM 5-FUrd

**Supplementary Figure 6:**
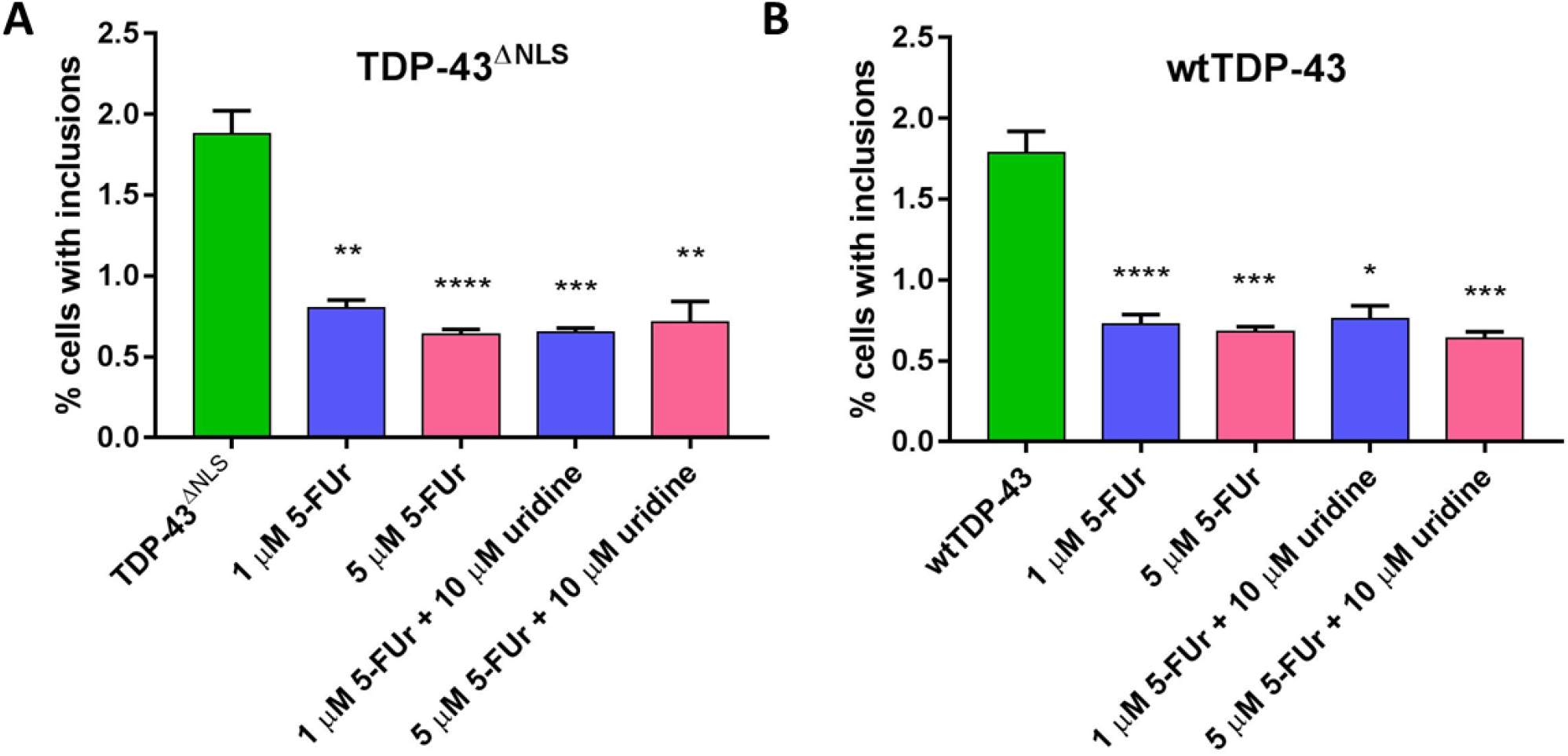
The presence of uridine does not decrease the effect of 5-FUrd to reduce SOD1 inclusion formation. HEK293FT cells were incubated with 5-FUrd and uridine 4-6 hours after co-transfection with TDP-43^ΔNLS^ and SOD-based reporter protein. Cells were collected 48 h post-transfection, treated with saponin and analyzed using flow cytometry for presence of aggregated reporter protein. Graphs represent the percentage of total cells with inclusion compared to transfected cells treated with vehicle control (green TDP-43^ΔNLS^ column). Statistical significance was determined using one-way ANOVA and Dunnett’s test for multiple comparisons (* p <0.05; ** p < 0.01; *** p < 0.001; **** p < 0.0001).

## MATERIALS AND METHODS

### Animal Ethics Statement

Use of zebrafish for this study was approved by the Animal Care and Use Committee: BioSciences at the University of Alberta under protocol AUP00000077, under the auspices of the Canadian Council on Animal Care. Adult zebrafish were maintained and bred according to standard procedures, including housing in brackish water (1250 ± 50 μS) at 28.5 °C.

### Materials

Brain samples were obtained from The Netherlands Brain Bank, Netherlands Institute for Neuroscience (Amsterdam, The Netherlands). Both donors had a clinical diagnosis of sporadic ALS with TDP-pathology. They gave written informed consent for CNS autopsy and the use of the material and clinical information for research purposes.

SOD1^G85R^-GFP plasmid was the kind gift from Professor Elizabeth Fisher (University College London, UK) (64) (Addgene: 26410), and a plasmid containing human wild-type TDP-43-FLAG was kindly provided by Dr. M. Urushitani (Shiga University of Medical Science, Japan). A plasmid for expression and purification of the N-terminal domain of TDP-43 (residues 1-80) (65) was generously provided by Dr. Nicolas L. Fawzi (Brown University, USA).

Other materials were purchased from the following companies: Human embryonic kidney cells (HEK293FT and HEK293; ATCC, Manassas, VA), L-glutamine (#25030081,Thermo Fisher, USA), Penicillin-Streptomycin (#15140122, Thermo Fisher, USA), Lipofectamine LTX (#15338100, Thermo Fisher), JetPrime transfection reagent (Polyplus, France), Expand High Fidelity PCR System (#11759078001, Roche, Switzerland) rabbit anti HA-tag antibody (#ab9110, Abcam, USA), chicken anti-HA tag antibody (#ab9111, Abcam, USA), mouse anti-TDP-43 antibody (#60019-2-Ig, Proteintech, USA), mouse anti-misfolded SOD1 monoclonal antibody 3H1 generated by Cashman lab (11), rabbit anti-TDP-43 C-Terminal antibody (#12892-1AP, Proteintech, USA), rabbit anti-SOD1-100 antibody (#ADI-SOD-100, Enzo Life Sciences), mouse anti-HA antibody (#ab130275, Abcam), rabbit anti-phospho S409/410 TDP43 antibody (Tip-PTD-Mo1, Cosmo Bio, Japan), Bis-benzimide H33342 trihydrochloride (#H3570, Thermo Fisher, Canada), Fluoromount-G (#00-4958-02, SouthernBiotech, USA). Secondary antibodies conjugated to fluorophores that were used for immunocytochemistry (Alexa-Fluor) were raised in goat and purchased from Thermo Fisher, Canada. Trp68 rabbit polyclonal antibodies were custom generated and affinity purified against the TDP-43 amino acid sequence _65_DAG<u>W</u>GNL_71_ by GenScript (Piscataway, NJ).

For immunohistochemistry, we used a kit containing both the HRP labelled goat anti-rabbit/mouse secondary antibody and the 3,3-Diaminobenzidine, called REAL EnVision Detection system (#K5007, Dako, Denmark), we used a universal antibody dilution buffer whenever the buffer composition is not specified (#U3510-100ML, Sigma, USA), and for mounting, we used Quick-D Mounting medium (#7281, Klinipath, Netherlands). Mouse anti-myc 488 conjugated was purchased from Millipore (#16-308). Superfrost plus tissue slides (Menzel-Gläser, Germany), FastDigest NotI restriction enzyme (#FD0593, Thermo Fisher, Canada), Ambion mMESSAGE SP6 transcription kit (#AM1340, Thermo Fisher, Canada), Clear Frozen Section Compound (#95057-838, VWR), Superfrost Plus microscope slides (#12-550-15, Fisher Scientific).

Microscope objectives: 63× objective on the Leica TCS SP8 microscope (#506350, Leica, Germany), A-Plan 10× objective, (#421040-9900-000, Carl Zeiss AG, Germany) equipped with an AxioCam HighRes camera (Carl Zeiss AG, Germany).

For Electrophoresis and Immunoblotting, we purchased pre-cast NativePAGE™ 4–12% Bis-Tris gels and NuPAGE™ 4–12% Bis-Tris Protein Gels, 1.0 mm (#NP0303BOX & NP0321BOX respectively, Thermo Fisher Scientific, USA), NativePAGE™ Sample Buffer (#BN2003, Thermo Fisher Scientific, USA), NuPAGE™ LDS Sample Buffer (# NP0007, Thermo Fisher Scientific, USA), 0.45 μm PVDF Transfer membrane (#88518, Thermo Fisher Scientific, USA), XCell II™ Blot Module (#EI9051,Thermo Fisher Scientific, USA). Amersham ECL HRP-conjugated secondary donkey anti-rabbit antibody (#NA93V, GE Healthcare Life Sciences, USA). SuperSignal™ West Femto Maximum Sensitivity Substrate (34094, Thermo Fisher Scientific, USA) and the ChemiDoc MP Imaging system (Biorad,USA) were used for detection.

For Size Exclusion Chromatography, we used Superdex 75 (10/300) HPLC column (#17517401, GE Healthcare Life Sciences, USA)

### Transfections

HEK293FT and HEK293 cells were cultured in complete Dulbecco’s Modified Eagle Medium (DMEM) containing 10% FBS, 10 U/ml penicillin, 10 U/ml streptomycin and 2 mM L-glutamine.

Systematic substitution of tryptophan residues in wtTDP-43 and TDP-43^ΔNLS^ was performed using site-directed mutagenesis. All of our TDP-43 constructs are HA-tagged or Flag-tagged to distinguish between endogenous and exogenous protein.

Pre-plated cells were co-transfected with the reporter protein (SOD1^G85R^-GFP), and one of the TDP-43 constructs at a ratio of 1:5, respectively, using Lipofectamine LTX according to manufacturer’s instructions. For the quantification of TDP-43-induced SOD1 misfolding, cells were co-transfected with HA-tagged wild-type TDP-43 and scFv antibodies.

For measurement of TDP-43-induced SOD1 misfolding, plasmid containing human wild type TDP-43-Flag was co-transfected with the pscFv9 plasmid (29) encoding for VH1Vk9, VH7Vk9 anti-RRM1 scFv antibodies, D1.3 scFv anti-chicken lysozyme scFv antibody, or pscFv9 empty plasmid. JetPrime transfection reagent was used according to manufacturer’s instructions. Mouse mAb 3H1 immunocytochemistry was used to quantify the conversion of wild-type endogenous human SOD1 to a misfolded form.

Cells were then incubated for 48 h in a 37 °C humidified incubator supplemented with 5 % CO_2_. In some instances, cells were treated with 5’-fluorouridine (5-FUrd). The drug was dissolved in DMEM media under sterile conditions and added at the indicated doses 4 h following the transfection of cells (13).

### Quantification of SOD1 Inclusions using flow cytometry

The abundance of cells with induced aggregation of SOD1^G85R^-GFP reporter protein was determined using flow cytometry (13, 20). In order to identify only those cells that not only express SOD1^G85R^-GFP, but where this report protein is indeed aggregated, we permeabilized the cells using 0.03% saponin in ice-cold PBS for 10 minutes to allow soluble SOD1^G85R^-GFP to leak out of cells (13, 20). Cells were then washed once using cold PBS and were stored on ice until analysis on LSRII (BD Biosciences, USA). Data analysis was performed using FlowJo.

### Immunocytochemistry

HEK293FT cells were seeded at a density of 6.3 × 10^5^ cells/cm^2^ on Poly-D-Lysine coated glass coverslips (#1.5 thickness) in a 24 well plate prior to transfection. 48 h post-transfection, cells were washed twice with ice-cold PBS and fixed in 4% paraformaldehyde (in PBS, pH 7.4) for 15 min at room temperature. Fixed cells were washed once with PBS, permeabilized for 10 min using PBSTx (0.3% Triton X-100 in PBS), and blocked for 30 min with blocking buffer (10% normal goat serum in PBS, filtered). Cells were incubated with primary antibodies diluted in incubation buffer (10% normal goat serum in PBS, filtered) to the following concentrations: 1 μg/ml for both chicken and rabbit anti-HA antibodies, 0.5 μg/ml rabbit anti-Trp68 for 1 h at room temperature. Cells were washed twice in PBS, and incubated with the appropriate secondary antibody (Alexa-fluor 488 anti-mouse, Alexa-fluor 568 anti-chicken, Alexa-fluor 647 anti-rabbit) diluted 1:1000 in blocking solution for 1 h at room temperature in the dark. Cells were then washed with PBS, and DNA was counterstained using 2 μg/ml Hoechst 33342 for 5 min. Following two final washes the cells were mounted on a glass slide in a drop of Fluoromount-G. Confocal images were captured using Leica TCS SP8 microscope on the oil-immersion 63× objective, with a numerical aperture of 1.4 using the LAS-X software (Leica, Germany) at 2048 × 2048 pixel resolution. Images were acquired at the same settings, brightness and contrast were enhanced uniformly across treatment groups for clarity where indicated.

#### Imaging and quantification of SOD1 inclusions using microscopy

Time-lapse live-cell imaging was carried out on HEK293FT cells cultured in 8-well microscope chambers. Quantification of SOD1^G85R^-GFP reporter protein aggregation was performed as previously described (13). Briefly, the chamber was placed into a microscope-mounted incubation system to maintain ideal cell culture parameters (humidified, 5% CO_2_ at 37 °C). Image acquisition began 16 h post-transfection (to allow protein expression) using an A-Plan 10× objective with a numerical aperture of 0.25 mounted on an inverted Axio Observer Z1 microscope equipped with an AxioCam HighRes camera (Carl Zeiss AG, Germany) and a motorized stage. Images were acquired at 1024 × 1024 pixel resolution (3× 3 grid – stitched post-acquisition) every 30 min. Images were exported as high resolution JPG files.

Quantifying induced aggregation of SOD1^G85R^-GFP reporter protein utilized a custom algorithm developed in ImageJ that reliably measured inclusions based on the area that inclusions normalized to total expressed GFP (13, 20). Briefly, images are converted into 8-bit images, local background is subtracted via rolling-ball method, threshold is set based on background soluble reporter protein, and aggregate size is quantified (minimum size of 5 pixels to avoid noise; maximum size was set to one third of an average cell size). Total fluorescence area of all fluorescent reporter protein is also quantified. The percentage of aggregation is reported as a ratio of area of inclusions divided by total area of fluorescence. Aggregate abundance was reported as a percentage of the aggregate abundance observed at various experimental time-points or end-stage of the experiment when using Trp-containing construct. Statistical significance was determined using one-way ANOVA and Dunnett’s test for multiple comparisons, calculated in GraphPad Prism 7.

### Quantification of TDP-43-induced SOD1 misfolding

HEK293 cells were co-transfected with wild-type TDP-43 and scFv antibodies and stained as indicated above with 2 ug/mL 3H1 anti-misfolded SOD1 mouse mAb and 0.4 ug/mL anti-TDP43 C-Terminal antibodies. After incubation of the slides with the appropriate secondary antibodies, scFv antibodies were detected with incubation with mouse anti-myc 488 conjugated diluted 1:500 in PBS for 2 h.

Images were acquired using confocal microscope BX-61 Virtual Stage (Olympus) with a Z-stack and analyzed with ImageJ software. To quantify SOD1 misfolding, the total integrated density was measured for the 3H1 antibody signal in each picture after adjusting threshold to discard signal coming from the empty coverslip. Intensity was then normalized to the total number of cells defined as those positive for nuclear TDP-43 signal. Statistical analysis was a one-way ANOVA followed by Tukey’s multiple comparison test, performed in Prism 5.0 (GraphPad, La Jolla, CA, USA). A *p*-value lower than 0.05 was considered significant.

### Immunohistochemistry on paraffin embedded human tissue

Cervical spinal cord sections (8 μm thick) from formalin-fixed paraffin embedded issue were mounted on Superfrost plus tissue slides and dried overnight at 37 °C. Sections were deparaffinized and subsequently immersed in 0.3% H_2_O_2_ in methanol for 30 minutes to quench endogenous peroxidase activity.

The slides were subjected to heat pretreatment with Tris-EDTA and incubated overnight at 4 °C in 0.1 ug/mL rabbit anti-Trp68 diluted in a purchased antibody diluent. Omission of the primary antibodies served as a negative control, and a section was incubated with 1:8000 rabbit phospho-TDP43 antibody as a positive control. Secondary EnVisonTM HRP goat anti-rabbit/mouse antibody incubation was for 30 min at room temperature. 3,3-Diaminobenzidine from the same kit was used a chromogen for 10 minutes. Sections were counterstained with haematoxylin to visualize the nuclei of the cells, dehydrated and mounted using Quick-D mounting medium.

### Expression of human SOD1 and TDP-43 in zebrafish

The human SOD1 and TDP-43 genes, including versions with variations indicated, were cloned using Gateway^22^ recombination into the pCS2+ expression vector construct for mRNA synthesis (pCS2+.wtSOD1.pA, pCS2+.SOD1^Trp32S^.pA, pCS2+.HA-wtTDP-43.pA, pCS2+.HA-TDP-43^ΔNLS^.pA, pCS2+.HA-TDP-43^ΔNLS-Trpless^.pA, pCS2+.HA-TDP-43^W68,W113S^.pA, and pCS2+.HA-TDP-43^W68,W172S^.pA). mRNA production was done using FastDigest NotI restriction enzyme and the Ambion mMESSAGE SP6 transcription kit. mRNA was co-injected with 100 pg mCherry mRNA into 1-2-cell stage embryos from *mnx1:GFP* transgenics (ZFIN ID: ZDB-ALT-051025-4) crossed to wild type AB fish. Embryos were screened at 24 h post-fertilization (hpf) for mCherry fluorescence indicating successful injection, and any embryos with a disrupted body axis or other overt defects were excluded from analysis.

Effective dosages of SOD1 or TDP-43 mRNA were determined empirically, with previous publications as guides (14), to robustly induce a measurable phenotype above background (control mRNA) levels, but below maximum levels of axonopathy, and was established at 900 pg. To harmonize amount of mRNA injected between various groups, total mRNA dose was made constant by top-up with innocuous Tol2 transposase mRNA, for a total 1900 pg mRNA per embryo. A control group was utilized to account for injection of exogenous mRNA; the mRNA control group dose consisted of 1800 pg of Tol2 transposase mRNA and 100 pg mCherry mRNA.

### Assessment of axonopathy

Embryos were raised to 36 hpf, fixed briefly in 4% formaldehyde, and assessed for axonopathy via GFP expression in the primary motor neurons by an observer blinded to the treatments. Dysmorphic embryos were not assessed. The axonopathy phenotype is an established robust assay for abnormal development/maintenance of the primary motor neurons caused by overexpression of wild-type and mutant SOD1, TDP-43, and other genes associated with human neuromuscular disease (22). Across this variety of diseases, intensity of the axonopathy phenotype is strongly correlated with disease severity. Briefly, primary motor axons were scored as abnormal if they exhibited branching dorsal of the notochord’s ventral boundary; total counts of abnormal axons were recorded per embryo (Fig. 2 A). Statistical analysis was performed using Kruskall-Wallis ANOVA with post-hoc Mann Whitney pairwise comparisons in Stata/SE 14.1 for Mac (2015, StataCorp). In some instances, muscle fibers were visualized by staining with Alexa Fluor-555-tagged phalloidin for 1 h. Embryos were subsequently mounted on slides using 1.5% low-melting point agarose and imaged on a Zeiss LSM 700 confocal laser microscope with Zen 2010 software (Carl Zeiss Imaging).

### Drug-based rescue using 5’-fluorouridine in zebrafish

The effective dosage of 5-fluorouridine (5-FUrd) was determined by establishing a dose-response curve for survival and axonopathy in zebrafish. Embryo media was replaced with media containing either drug or vehicle control when embryos reached 12 hpf. 5-FUrd treatment media contained 1.5 μM 5-FUrd, 5 μM uridine, and 0.2% DMSO, whereas vehicle control media contained 5 μM uridine and 0.2% DMSO. Embryos were kept in drug media until fixation in 4% formaldehyde at 36 hpf.

### Zebrafish Immunohistochemistry

For labeling to detect human SOD1 and HA-TDP-43^ΔNLS^ in zebrafish, embryos (without GFP transgenes) were fixed at 30 hpf and cryopreserved as previously described using step-wise sucrose/0.1 M PO4 washes and freezing in sucrose/PO4 mixed with Clear Frozen Section Compound overnight at −80 °C (22). 10 μm cryosections were mounted on Superfrost Plus microscope slides, allowed to air-dry, and frozen at −80 °C overnight. Slides were then thawed, incubated in 10% normal goat serum/PBSTw, and incubated in primary antibodies overnight: 1:500 rabbit anti-SOD1-100 (Enzo Life Sciences, ADI-SOD-100) and 1:100 mouse anti-HA (Abcam, ab130275). After washes with PBSTw, slides were incubated in Alexa-fluor 488 anti-rabbit and Alexa-fluor 647 anti-mouse antibodies overnight. Following final washes, slides were imaged on a Zeiss LSM 700 confocal laser microscope with Zen 2010 software (Carl Zeiss Imaging).

### Purification of TDP43 NTD

Protein was expressed in Escherichia coli and purified as described previously (65). Purified TDP-43 NTD was dialyzed against PBS, concentrated to 0.5 mg/ml and stored at −80 °C.

### Electrophoresis using SDS/Native gels and Immunoblotting

Native-PAGE was carried out using the Novex Bis-Tris system according to the manufacturer’s specifications. Protein samples were mixed with the NativePAGE™ Sample Buffer. Pre-cast NativePAGE™ 4–12% Bis-Tris gel was run at 4 °C at 150 V constant for 60 min, then at 250 V for the 30 min.

SDS-PAGE was carried out using the Novex Bis-Tris system according to the manufacturer’s specifications. Protein samples were mixed with the NuPAGE™ LDS Sample Buffer. Pre-cast NuPAGE™ 4–12% Bis-Tris gel was run at RT at 200 V constant for 35 min.

Proteins were blotted onto 0.45μm PVDF membranes using the XCell II Blot Module following the manufacturer’s protocol. Blots were blocked in 5% milk powder in 0.0.2% Tween 20 Tris Buffer Saline, and then were incubated with 0.5 μg/mL Trp68 antibody overnight at 4°C. For detection on the ChemiDoc MP, a donkey anti-Rabbit IgG HRP-labelled secondary antibodies was used. The SuperSignal West Femto substrate was used according to the manufacturer’s instructions.

### Size Exclusion Chromatography

Analytical gel-filtration of TDP43 NTD forms was performed using high performance liquid chromatography on Superdex 75 (10/300) HPLC column. TDP-43 NTD (0.5 mg/ml) was denatured by incubation in 6 M Guanidine-HCl, 50 mM Tris-HCl buffer (pH 7.5.), 150 mM NaCl, 20 mM DTT, for 10 min at 37 °C. The obtained protein sample (100 μl) was loaded onto the column pre-equilibrated with the same buffer and eluted at 0.5 ml/min. Native form was loaded onto the column pre-equilibrated with the 1x PBS buffer containing 5 mM DTT and eluted at 0.5 ml/min.

### Computational Protein Modelling

Representative structures were obtained as the top-ranked structures using 3D-Jury ranking with MaxSub comparison between structures and threshold RMSD of 2 (66). The MaxCluster program was used to obtain the 3D-Jury ranking (67). Native conformational ensembles of the N-terminal domain and RRM1 domain of TDP43 were obtained from equilibrium molecular dynamics simulations using GROMACS 4.5 and 5.0 respectively (68). Apo structures without nucleic acid or other ligand were used to calculate the SASA of Tryptophans in a non-redundant database of 27,015 structures taken from the PDB, as described in (27).

### Statistical Analysis

Statistics were performed as described above in figure legends.

